# Informative auditory cues enhance motor sequence learning

**DOI:** 10.1101/2024.10.07.617139

**Authors:** Li-Ann Leow, An Nguyen, Emily Corti, Welber Marinovic

## Abstract

Motor sequence learning, or the ability to learn and remember sequences of actions, such as the sequence of actions required to tie one’s shoelaces, is ubiquitous to everyday life. Contemporary research on motor sequence learning has been largely unimodal, ignoring the possibility that our nervous system might benefit from sensory inputs from multiple modalities. In this study, we investigated the properties of motor sequence learning in response to auditory-visual stimuli. We found that sequence learning with auditory-visual stimuli showed a hallmark feature of traditional unimodal sequence learning tasks: sensitivity to stimulus timing, where lengthier interstimulus intervals of 500 ms improved sequence learning compared to briefer interstimulus intervals of 200 ms. Consistent with previous findings, we also found that auditory-visual stimuli improved learning compared to a unimodal visual-only condition. Furthermore, the informativeness of the auditory stimuli was important, as auditory stimuli which predicted the location of visual cues improved sequence learning compared to uninformative auditory stimuli which did not predict the location of the visual cues. Our findings suggest a potential utility of leveraging audio-visual stimuli in sequence learning interventions to enhance skill acquisition in education and rehabilitation contexts.

## 1. Introduction

Organizing actions, events, words, memories and thoughts into sequences is a critical part of everyday behavior (Ashe et al, 2006). The ability to learn sequences of goal directed actions is ubiquitous and necessary in actions ranging from typing this sentence using a keyboard to complex tasks like language and musical performance. Whilst the bulk of research on sequence learning have focused primarily on learning sequences based on visual inputs (Nissen & Bullemer, 1987; Cohen *et al*., 1990; Curran & Keele, 1993; Hoffmann & Koch, 1997), recent studies have sought to understand how multisensory stimuli can affect sequence learning (Silva *et al*., 2017; Luan *et al*., 2021; Han *et al*., 2023). This is in line with increasing recognition that we live in a multisensory world and that our nervous system likely capitalizes on sensory inputs from multiple modalities (Seitz *et al*., 2006a; Shams & Seitz, 2008). For example, multisensory information can enhance a range of learning processes, including visual learning (Seitz *et al*., 2006b; Kim *et al*., 2008), language acquisition (Vigliocco *et al*., 2014), and mathematical skills (Baker & Jordan, 2015)

Multisensory information can be implemented as a form of *feedback* (Stocker *et al*., 2003; Stocker & Hoffmann, 2004; van Vugt & Tillmann, 2015; Luan *et al*., 2021; Robinson & Parker, 2021), or simply as a form of *stimuli* without associated information about the action (Silva *et al*., 2017; Han *et al*., 2023). Both can improve performance, although the pattern of performance improvement seems to differ depending on the *informativeness* of the multisensory information (Stocker *et al*., 2003; Stocker & Hoffmann, 2004; Luan *et al*., 2021), for example, where auditory stimuli provides information about the location of visual stimuli. Similarly, improved performance might also depend on the *timing* of the multisensory information, as well as the explicit or implicit nature of the learning task. For example, Silva *et al*. (2017) investigated how different forms of auditory information presented synchronously with visual stimuli (informative tones that carry information about the order of keypresses, random tones, single tones, and no tones) affected performance in an implicit sequence learning task. They found that whilst auditory information had no effect on performance, auditory stimuli congruent with visual stimuli tended to increase the development of explicit knowledge of the sequence to be learnt, which correlated with the magnitude of learning. In a similar vein, under conditions which facilitate explicit learning (i.e., lengthy movement preparation times), providing auditory cues synchronously with visual cues seem to enhance performance compared to visual-only training: in a version of the sequence learning task where visual stimuli move across the screen over 1500ms, auditory stimuli presented in synchrony with the start of the visual stimuli improves sequence learning (Han *et al*., 2023). This differs from typical sequence learning tasks where participants respond to briefly presented visual stimuli, requiring actions with minimal preparation.

The timing of stimuli seems to play a critical role in determining the extent of explicit sequence learning. In turn, the extent of explicit sequence learning appears to modulate how effectively multisensory information enhances performance. In Silva et al.’s (2017) study, they used brief intervals (200 ms) between response execution and the presentation of the next stimulus in the sequence. These brief response-stimulus intervals are known to impair sequence learning (Destrebecqz & Cleeremans, 2001), and may have impacted critical error-monitoring mechanisms active immediately after each response, which influences the extent of learning (e.g., Hadipour-Niktarash *et al*., 2007; Huang & Shadmehr, 2007). Such brief post-response intervals might thus have limited observable benefits from congruent audio-visual stimuli in the Silva et al., 2017 study.

To the best of our knowledge, the timing of multisensory information and the congruence of multisensory information has not yet been systematically tested within the context of multisensory sequence learning. In the present study, we sought to investigate the effect of the timing of multisensory information and the congruence of multisensory information on sequence learning task performance. In our first study, in a within-participants’ design, we manipulated the interval between the execution of a response and the presentation of the next visual stimulus in the sequence, whilst informative auditory stimuli (i.e., each tone corresponded to a specific visual stimulus) were presented in synchrony with each visual stimulus. We predicted that participants would show faster learning when the sequence to be learned was presented more slowly: 500 ms vs. 200 ms. Our second study used a between-subjects design to examine how the informativeness of auditory stimuli altered sequence learning. Auditory stimuli were presented in synchrony with visual stimuli, and were either ***informative*** (each tone corresponded to each visual stimulus), ***uninformative*** (the same tone for all visual stimuli, providing no informative auditory differentiation), or ***random*** (randomized tones for all visual stimuli, precluding any predictable association between the auditory and visual stimuli). This was compared to a control condition with no auditory stimuli (no tones). We hypothesized that informative audio-visual stimuli would lead to a faster learning in comparison to the other groups. In our final study, we conducted a direct comparison using a repeated measures design, contrasting performance differences during training when sequences were paired with informative versus uninformative stimuli. Here, we predicted that informative stimuli would result in faster sequence learning.

## 2. Method

### 2.1. Participants

A total of 201 participants were recruited from the undergraduate psychology program at Curtin School of Population Health. They volunteered to participate in exchange for course credit. For study 1, we collected data from 32 participants aged between 18 and 48 years (Mean = 22.4, SD = 6.6, 28 female). For study 2, data from 137 participants aged between 17 and 48 years (Mean = 21.8, SD = 5.7, 91 female) were collected. Participants were randomly allocated to one of four experimental groups in this study. For study 3, we gathered data from 32 participants aged between 18 and 50 years (Mean = 21.7, SD = 6.4, 19 female). All participants had normal or corrected-to-normal vision, normal hearing, and had no known or diagnosed neurological conditions that could have affected their performance in the task. Participants provided signed consent prior to participation in the experiment. This study was approved by the Curtin Human Research Ethics Committee (HRE2018-0257).

### 2.2. Sample size

We did not conduct a formal sample size analysis. Instead, we aimed to collect approximately 32 participants per group for each study. This sample size exceeds the sample size of 20 participants per group used by Silva *et al*. (2017), who similarly presented auditory cues as stimuli in a motor sequence learning task. We reasoned that this larger sample size would align with typical effect sizes observed in such research and ensure adequate power to detect effects of interest.

### 2.3. Apparatus

Visual and auditory stimuli were presented using Inquisit 6 software. Participants were seated in front of a computer monitor (1920 x 1200, 60Hz, 24-inch) at a distance where they could comfortably reach a standard QWERTY keyboard placed between them and the monitor screen. Sounds were presented through stereo headphones (Corsair HS50) at a fixed intensity of 65dBa and had a duration of 100 ms.

### 2.4. Procedures

Across all studies, participants were instructed to learn sequences of keypresses by responding to four grey squares on a computer screen that turned red according to a fixed sequence. Participants placed their left middle and index fingers on the ‘V’ and ‘B’ keys, respectively, and their right index and middle fingers on the ‘N’ and ‘M’ keys, respectively. Four grey squares were located horizontally on the computer screen and mapped to these keys (V, B, N, and M) moving left to right. Stimuli were either unimodal or bimodal, depending on the study design. In all trials (unimodal or bimodal), the grey square “flashed”, (i.e., it changed its color to red). In bimodal trials, as the grey square changed color, a pure tone (C4, D4, E4, or F4) simultaneously played through the headset. These pure tones C4 (261.63 Hz), D4 (293.66 Hz), E4 (329.63 Hz), and F4 (349.23 Hz) varied depending on the task manipulation in each study, described later. Participants responded as quickly and accurately as possible to the stimuli, by pressing the key that corresponded with the square that changed color. Before participants started the experiment, they performed ordered practice trials (‘1,2,3,4,1,2,3,4’) as well as random trials (i.e., trials where the stimuli appeared in random order) to familiarize themselves with the task.

In Study 1, we employed a within-subjects design, manipulating the response-stimulus interval (i.e., the interval between each response and the subsequent stimulus)(Willingham *et al*., 1997), which was either 200 ms or 500 ms. Across all trials, stimuli were bimodal (i.e., auditory tones were paired with each color change), and each square location was mapped 1:1 to each auditory tone (this is thereafter termed Bimodal Informative mapping). Participants were required to learn two distinct sequences of keypresses, with each sequence comprising 10 items (‘1,3,1,4,3,2,4,2,3,1’, and ‘3,2,4,1,3,3,4,2,1,2’). They were explicitly instructed to learn both sequences equally. Each sequence used either the 200 ms or the 500 ms response-stimulus interval. Participants performed each sequence 10 times consecutively, resulting in a block of 100 keypresses. After completing a block with one sequence, they switched to the other sequence, with instructions on screen to warn them about the switches. This process was repeated until each participant completed a total of 1,000 keypresses, evenly divided between the two sequences (500 keypresses per sequence), across alternating blocks of 100 trials each. The presentation order of sequences as well as the response-stimulus interval was counterbalanced across participants.

Study 2 used a between-subjects design to investigate how different auditory stimuli mappings altered sequence learning. Here, all participants were required to learn only one sequence of keypresses, which was fixed for all participants (‘3,1,3,1,4,2,4,4,1’), and the response-stimulus interval was held constant at 500ms across all stimuli across all participants. Participants were randomly assigned into four distinct groups based on the type of stimuli they were presented with during the task: the Unimodal group was exposed only to the visual flashes without tones; the Bimodal Informative group received synced visual and auditory stimuli with a consistent mapping between the tones and the flashes; the Bimodal Uninformative group was presented with a single, repetitive tone alongside the visual flashes, providing no informative auditory differentiation; and the Bimodal Random group received a randomized tones with the fixed sequences of flashes, precluding any predictable association between the auditory and visual stimuli. As there was only one sequence to be learned, participants performed only 500 trials, divided into 5 blocks of 100 trials each.

Study 3 used a within-subjects design to further investigate how the informativeness of the auditory mapping altered sequence learning. Study 3 design was identical to Study 1, except that it changed the auditory mapping for one sequence to uninformative (where each of the four squares was associated with a single tone, thereafter termed the Bimodal Uninformative condition), whereas the other sequence was informative (each of the four squares were associated with a different tone, i.e., the Bimodal Informative condition). The response-stimulus interval was held constant at 500ms across all stimuli. Participants learned the two sequences (‘2,4,1,1,3,4,2,4,3,1’ and ‘1,4,3,1,2,2,4,3,4,1’). The order of sequence presentation as well as the mapping condition was counterbalanced across participants.

**Figure 1.**
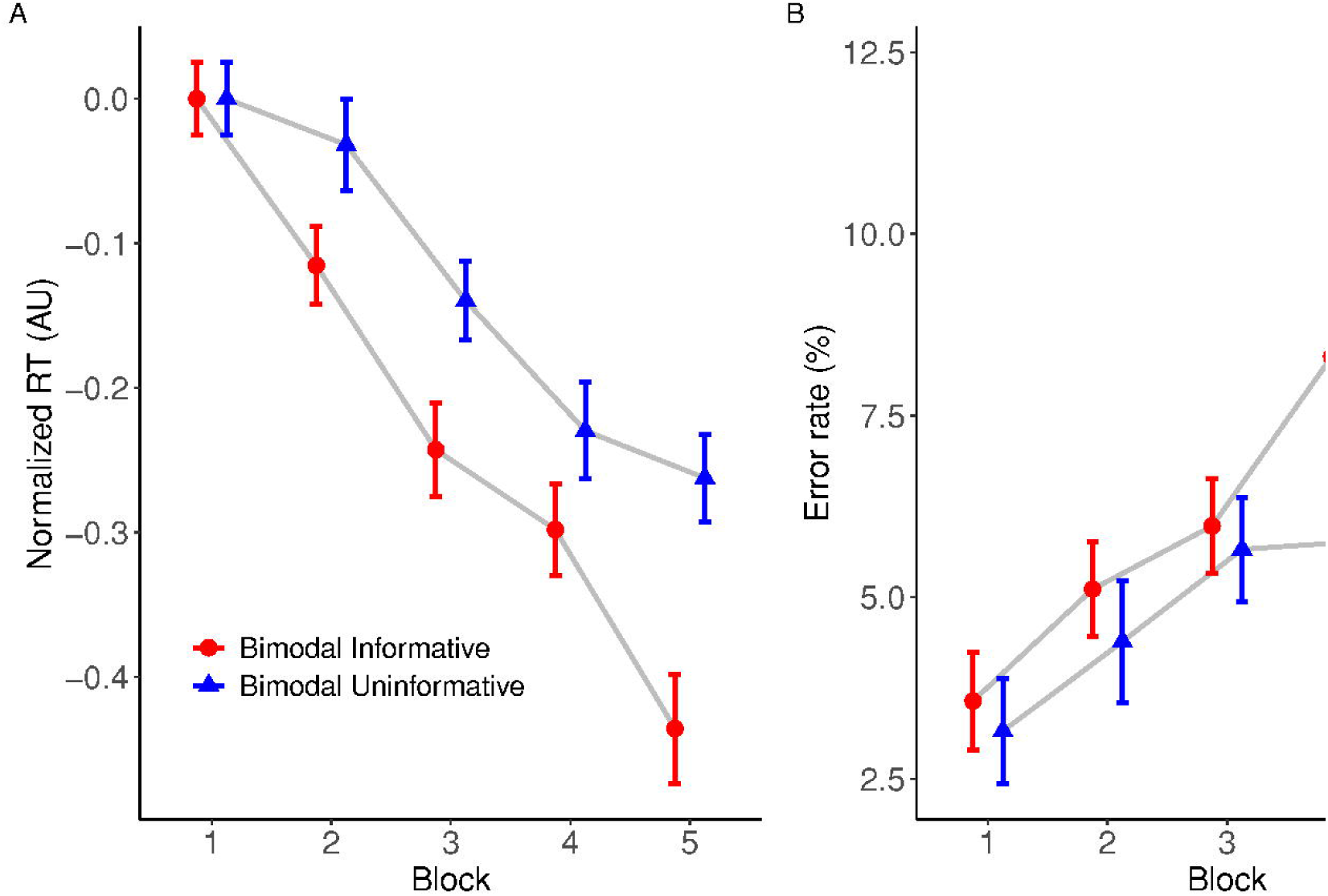
Experimental manipulations used in Study 1 (top) and Study 2&3 (bottom).

### 2.5. Data analysis

Statistical analyses were performed using R Statistics (version 4.3.2) and R Studio (Build 404). Our two main dependent variables of interest across all studies were normalized reaction time (RT) and error rates. Normalized RTs were obtained using the following equation 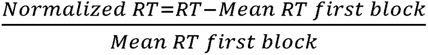 (Perez *et al*., 2007). For the normalized RT analysis, we discarded trials where participants made errors or when the response took longer than 1 second to occur. Therefore, error trials did not contribute to the estimation of RT effects. The percentage of discarded trials due to errors are as follows: study 1: 7.01%, study 2: 4.51%, study 3: 5.74%. We also discarded trials with RTs longer than 1000 ms, as follows: study 1: 2.50%, study 2, 1.89%, study 3: 1.78%.

For the analysis of error rates, only trials where participants took longer than 1 second were discarded but incorrect keypresses were necessary for the estimation of error rates using a logistic regression model. Following recommendations for the robust data analysis of repeated measures designs, all retained individual trials were included in our analysis of the linear and generalized mixed-effects models (Hedeker & Gibbons, 2006; Barr *et al*., 2013; Yu *et al*., 2022). Our analysis approach combined both frequentist and Bayesian methods, utilizing their complementary strengths to provide a more robust understanding of our data, in line with the recommendations made by Flores *et al*. (2022). We initially fitted a series of linear mixed models (LMMs) to our normalized RT data using the ‘lmer’ function from the ‘lmerTest’ package (Kuznetsova *et al*., 2015). Four different LMMs were compared: one with a linear term for Block and Condition, and three incorporating polynomial terms of different degrees for Block (2^nd^, 3^rd^, and 4^th^). These models were compared using the ‘anova’ function from the ‘stats’ package to identify the model that best explained the variation in normalized RT across blocks. For the normalized RT, the anova function was also used to extract the key metrics for the best model (degrees of freedom, F and p values). For the error rate analysis, we employed generalized linear mixed model using a logistic function and extract statistics from these models using the ‘Anova’ (type = 3) function from the ‘car’ package. To corroborate the frequentist analyses, we also employed Bayesian linear mixed models, providing a probability-based interpretation of our data. A series of models were compared using the ‘bayesfactor_models’ function from the bayestestR package (Makowski *et al*., 2019), including models with main effects and interactions, as well as a null model for baseline comparison. The inclusion of each term in the models was assessed using the Inclusion Bayes Factor (BF _inclusion_), calculated with the ‘bayesfactor_inclusion’ function from the ‘BayesFactor’ package (Morey *et al*., 2015). For all models, Condition (Study 1 and 3) or Group (Study 2) were entered into the models as a fixed factor, with Blocks entered as a numeric covariate, and participant ID as a random effect. All models reported converged successfully.

For tests involving multiple pairwise comparisons within or across experimental blocks, we applied the Tukey correction to control for Type I errors in our frequentist analyses. In contrast, for studies where only a single pairwise comparison was conducted within each block, no correction was employed. We only performed these follow up tests if the effects were supported by both the frequentist and Bayesian approaches. To obtain robust follow-up estimates of pairwise differences within the Bayesian framework, we implemented Bayesian linear models using the anovaBF function from the ‘BayesFactor’ package and extracted 95% credible intervals from the posterior distribution (using 50,000 iterations). Because these credible intervals were derived directly from a joint posterior distribution, our Bayesian approach inherently accounted for multiplicity by integrating uncertainty across comparisons, removing the need for additional post hoc adjustments (see Berry & Hochberg, 1999). Credible intervals are presented along with their respective Posterior Probability of Direction (PD) of effects, which represents the proportion of the posterior distribution that supports the estimated direction of the effect. A PD close to 1 or 0 suggests strong consistency in the effect’s direction, whereas values near 0.5 indicate uncertainty.

## 3. Results

### 3.1. Study 1: reaction time

The interstimulus interval influenced training-related reductions in normalized reaction time across training blocks (see Figure 2). The model including a third-degree polynomial resulted in the lowest AIC and BIC values, suggesting it was the most appropriate to explain our data. The ANOVA for this model revealed a statistically significant main effect of ‘Block’ (F_(3, 28914)_ = 759.89, p < 0.0001), reflecting reductions in reaction times as the blocks progressed. The main effect of ‘Condition’ failed to reach statistical significance (F_(1, 28914)_ = 1.58, p = 0.208). The interaction between ‘Block’ and ‘Condition’ was statistically reliable (F_(3, 28914)_ = 14.64, p < 0.0001), suggesting that changes in reaction time over blocks was affected by the timing condition. Corroborating these findings, an additional Bayesian analysis provided strong support for the inclusion of the factor ‘Block’ (BF _inclusion_= ∞) in the model as well as the interaction between ‘Block’ and ‘Condition’ (BF _inclusion_= 691). This analysis provided evidence against the inclusion of ‘Condition’ alone only (BF _inclusion_= 0.013), consistent with the non-significant effect provided by the ANOVA.

**Figure 2:**
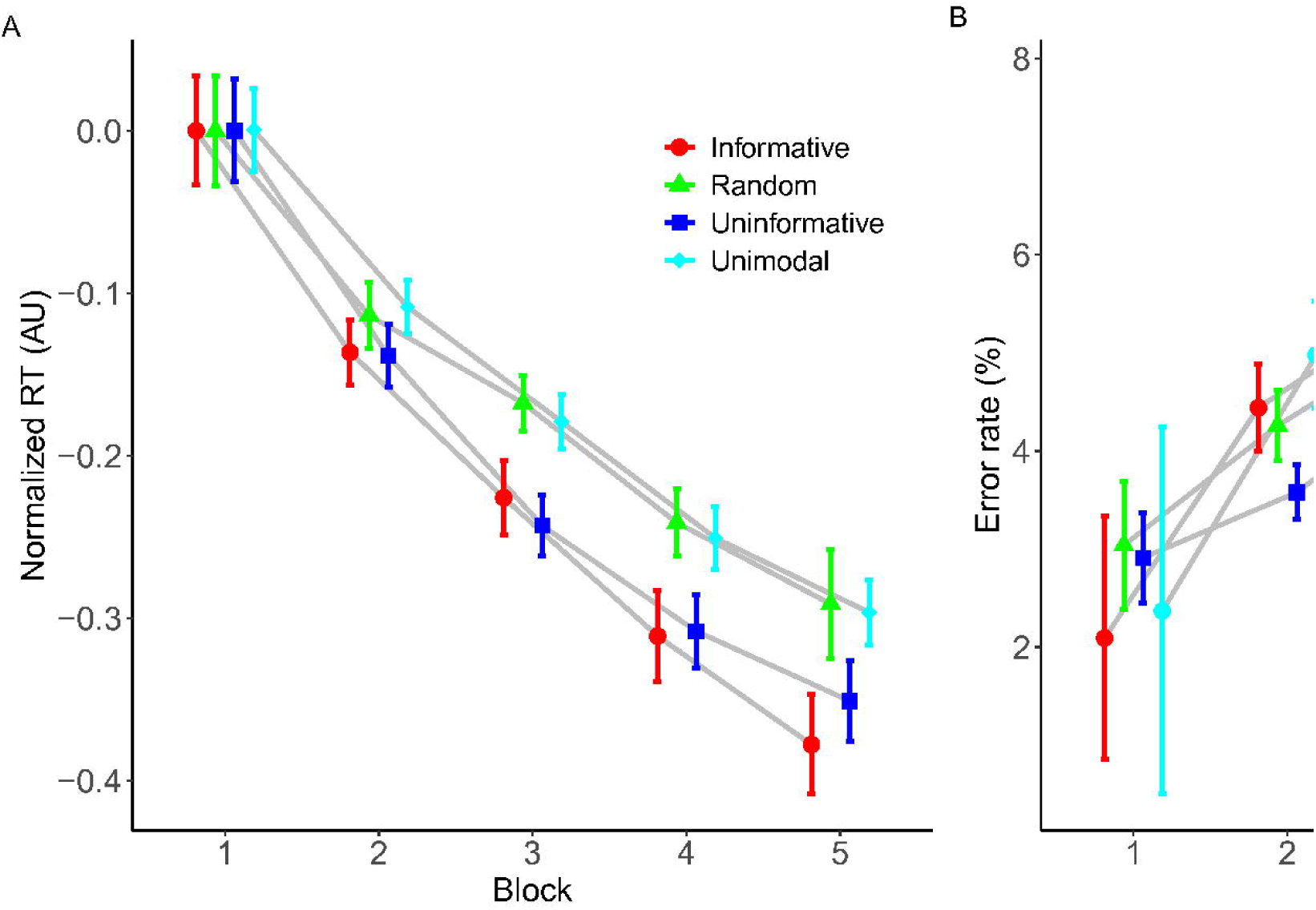
A- Average normalized reaction times for correct keypresses for the 200 ms and 500 ms interstimulus interval conditions across the five blocks. B - Average error rates for the 200 ms and 500 ms conditions across the five blocks. Error bars represent the within-participant standard errors (Morey, 2008).

A follow up on the interaction between ‘Block’ and ‘Condition’ revealed a dynamic pattern of results as shown in Figure 2A. Initially, the contrast between the 500 ms and 200 ms conditions in Block 1 showed no significant difference in reaction times (estimate = −0.000865, SE = 0.00961, z = −0.090, p = 0.99; Bayesian 95% CI: [−0.011, 0.024], PD = 0.76). However, a significant difference emerged at Block 2, where the 500 ms condition showed faster reaction times compared to the 200 ms condition (estimate = 0.0374, SE = 0.00857, z = 4.361, p < .0001; Bayesian 95% CI: [0.020, 0.057], PD = 0.99). In Block 3, there was no clear difference between conditions (estimate = −0.0104, SE = 0.00682, z = −1.522, p = 0.12; Bayesian 95% CI: [−0.013, 0.022], PD = 0.67). In Block 4, the 500 ms condition showed slower reaction times than the 200 ms condition (estimate = −0.0533, SE = 0.0086, z = −6.202, p < .0001; Bayesian 95% CI: [−0.072, −0.035], PD = 0.00). Finally, we detected no difference between conditions in Block 5 (estimate = −0.000698, SE = 0.00982, z = −0.071, p = 0.94; Bayesian 95% CI: [−0.014, 0.022], PD = 0.65). Therefore, while there was an initial advantage for the 500 ms Condition in Block 2, this advantage was reversed by Block 4. Both Conditions led to clear reductions in reaction time across Blocks and performance was by and large equivalent at the last Block of trials.

### 3.2. Study 1: error rate

Overall, the 200 ms interstimulus condition increased errors compared to the 500 ms condition (see Figure 2B). There was no significant difference between the linear and those including polynomials, so we report the results of the simple linear model. There was a statistically significant main effect of ‘Block’ (χ²= 31.57, df = 4, p < 0.0001), indicating that error rates increased over blocks (BF _inclusion_= 1.06e+17). Similarly, the main effect of ‘Condition’ was also significant (χ²= 27.35, df = 1, p < 0.0001), suggesting that the 200 ms condition induced more errors across blocks (BF _inclusion_= 2.37e+11). There was also a statistically significant interaction between ‘Block’ and ‘Condition’ (χ²= 5.30, df = 4, p = 0.021), reflecting some idiosyncrasies in the pattern of error changes across blocks depending on the condition. This interaction, however, was not corroborated by the Bayesian analysis (BF _inclusion_= 0.079). We did not conduct a post-hoc analysis on the interaction term for error rate as the Bayesian analysis did not support its inclusion in the model.

### 3.3. Study 2: reaction time

Training-related reductions in normalized reaction times differed depending on the stimulus group (Figure 3A). The model with a second-degree polynomial for ‘Block’ reliably outperformed the linear model, as indicated by lower AIC and BIC values and a statistically significant chi-square difference (χ² = 235.9, df = 4, p < 0.0001). Higher order polynomial models were not statistically superior to the second-degree polynomial model. The ANOVA of the second-degree polynomial model revealed a significant main effect of ‘Block’ (F_(2, 63874)_ = 4212.48, p < 0.0001), but the main effect of ‘Group’ failed to reach statistical significance (F_(3, 133)_ = 0.83, p = 0.479). The interaction between ‘Block’ and ‘Group’ was statistically significant (F_(6, 63874)_ = 20.57, p < 0.0001). Complementing these findings, the Bayesian analysis provided decisive support for the inclusion of the polynomial term of ‘Block’ (BF _inclusion_= ∞), and evidence against the inclusion of main effect of ‘Group’ (BF _inclusion_ = 2.13e-07). The interaction between ‘Group’ and ‘Block’ resulted in a very high BF (BF _inclusion_ = 2.33e+12), corroborating the frequentist results.

**Figure 3:**
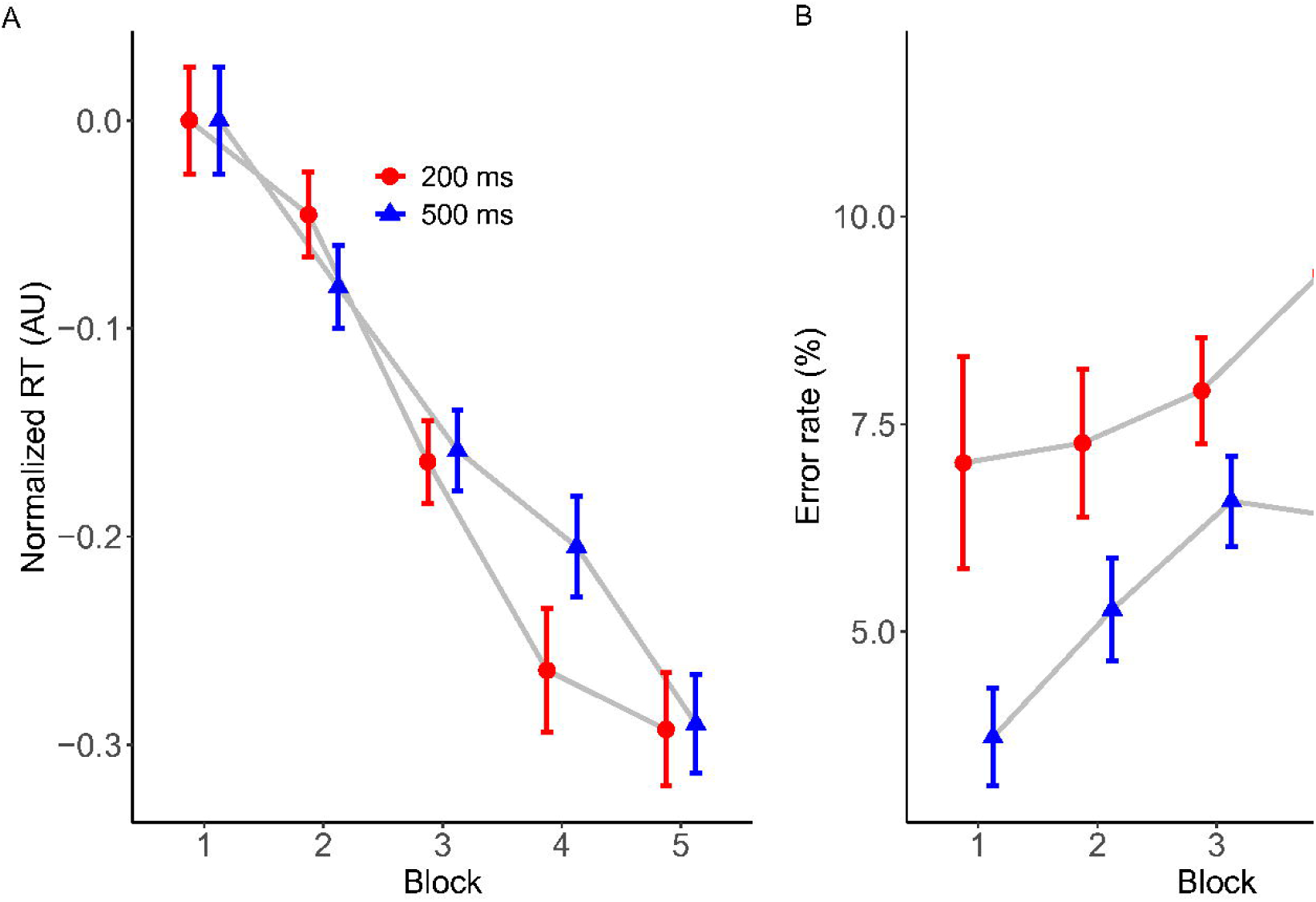
A- Average normalized reaction times for correct keypresses for the four Groups across the five blocks. B - Average error rates for the four Groups across the five blocks. Error bars represent the within-participant standard errors (Morey, 2008). Note that errors do not contribute to the normalized reaction time calculations.

While the pattern of results shown in Figure 3A suggests pairwise differences between conditions at specific blocks, frequentist post-hoc tests on the interaction between ‘Group’ and ‘Block’ did not identify significant pairwise differences across blocks after adjusting for multiple comparisons. However, the unadjusted p-values hinted at emerging differences in later blocks. Bayesian analyses provide additional support for these contrasts, particularly in Block 5. The group that received ‘Informative’ stimuli exhibited a likely advantage over the group trained with ‘Random’ stimuli (Bayesian 95% CI: [−0.054, −0.023], PD < 0.00), reinforcing the idea that informative cues may facilitate faster reaction times. Similarly, the contrast between the ‘Unimodal’ and ‘Informative’ groups was consistent with this trend (Bayesian 95% CI: [−0.042, −0.011], PD = 0.0002), suggesting that differences in reaction times became more pronounced in later blocks. While unadjusted frequentist p-values should be interpreted cautiously, the Bayesian results are consistent with our hypothesis that informative bimodal stimulation may benefit sequence learning.

### 3.4. Study 2: error rate

The results for error rate are shown in Figure 3B. The only model that successfully converged was the simple logistic linear model. This model revealed a statistically significant main effect of ‘Block’ (χ²= 80.511, df = 1, p < 0.0001), again showing that error rates increased as practice progressed. The main effect of ‘Group’ failed to reach statistical significance (χ²= 3.543, df = 3, p = 0.315). The interaction between ‘Block’ and ‘Group’ only approached statistical significance (χ²= 6.60, df = 3, p = 0.085). The Bayesian analysis provided evidence for the inclusion of ‘Block’ in the model (BF _inclusion_= 2.86e+45), with the main effect of ‘Group’ and the interaction both indicating against their inclusion in the model (Group: BF _inclusion_= 7.42e-08; Group x Block: BF _inclusion_= 1.48e-06).

### 3.5. Study 3: reaction time

Study 3 employed a within-subjects design to circumvent issues associated with individual differences in sequence learning (e.g., Stark-Inbar *et al*., 2017) that can contaminate effects of experimental manipulations using a between-subjects design. Within the same participants, we found that informative tones improved training-related reductions in reaction time compared to uninformative tones (see Figure 4A). The model comparison revealed that the model incorporating a fourth-degree polynomial for ‘Block’ significantly outperformed simpler models. This was evidenced by lower AIC and BIC values and a substantial chi-square difference (χ² = 22.787, df = 2, p < 0.0001). The ANOVA for this model found a reliable main effect of ‘Block’ (F_(4, 29552)_ = 834.25, p < 0.0001), reflecting reaction times reductions across the blocks. There was also a significant main effect of ‘Condition’ (F_(1, 29551)_ = 360.61, p < 0.0001). The ANOVA also revealed a statistically significant interaction between ‘Block’ and ‘Condition’ (F_(4, 29551)_ = 45.09, p < 0.0001). All these findings were corroborated by the Bayesian analysis (Block: BF _inclusion_= ∞; Condition: BF _inclusion_= 1.18e+75; Interaction: BF _inclusion_= 1.32e+30).

**Figure 4:**
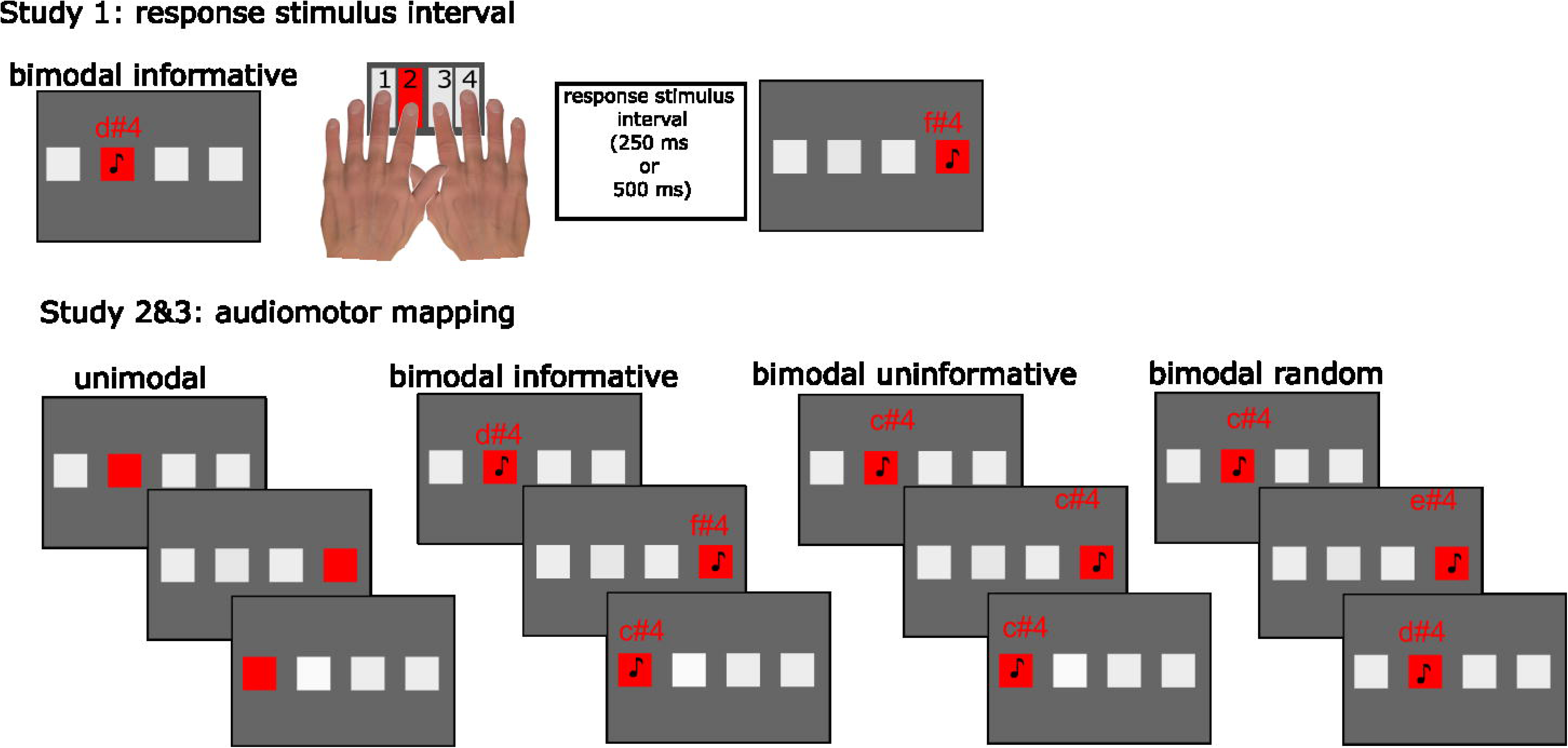
A- Average normalized reaction times for correct keypresses for the (bimodal) Informative and Uninformative conditions across the five blocks. B - Average error rates for the (bimodal) Informative and Uninformative conditions across the five blocks. Error bars represent the within-participant standard errors (Morey, 2008). Note that errors do not contribute to the normalized reaction time calculations.

As depicted in Figure 4A, a post-hoc follow up analysis on the interaction term found differences across all blocks but the first (Block 1: estimate = −0.0001, SE = 0.0097, z = −0.014, p = .98, Bayesian 95% CI: [0.064, 0.100], PD = 1.00; Block 2: estimate = −0.0831, SE = 0.0098, z = −8.47, p < .0001, Bayesian 95% CI: [−0.017, 0.019], PD = 0.52; Block 3: estimate = −0.104, SE = 0.0098, z = −10.505, p < .0001, Bayesian 95% CI: [−0.038, −0.002], PD = 0.016; Block 4: estimate = −0.0569, SE = 0.0099, z = −5.75, p < .0001, Bayesian 95% CI: [0.012, 0.049], PD = 0.99; Block 5: estimate = −0.182, SE = 0.0101, z = −18.05, p < .0001, Bayesian 95% CI: [−0.112, −0.075], PD = 0.00).

### 3.6. Study 3: error rate

Overall, error rates tended to increase with increasing training blocks (Figure 4B). There was no statistical difference between any of the models tested, so we report the simpler linear model here. The ANOVA showed a statistically significant main effect of ‘Block’ (χ² = 125.65, p < 0.0001) but not of ‘Condition’ (χ² = 0.17, p < 0.68). The Bayesian analysis corroborated the inclusion of the main effect ‘Block’ (BF _inclusion_= 7.79e+40), but contradicting the frequentist approach, it indicated that the effect of ‘Condition’ is important (BF _inclusion_= 369), which seems intuitive given the pattern of results shown in Figure 3B. The ANOVA also failed to find a significant interaction between ‘Block’ and ‘Condition’ (χ² = 2.18, p < 0.13), which is consistent with the Bayesian analysis (BF _inclusion_= 0.017). We did not conduct a follow up post-hoc test for this interaction.

## 4. Discussion

Here, we conducted a series of studies designed to address key issues regarding bimodal sequence learning. Recent studies have failed to convincingly demonstrate beneficial effects of bimodal sensory stimulation during learning when sensory stimuli were delivered in sync (Silva *et al*., 2017) in an implicit learning SRT task. We examined whether the interval between the response and the next stimulus in the sequence (i.e., the response-stimulus interval) could impact the rate of learning with bimodal stimuli. In our second study, following up on Silva and colleagues’ design (Silva *et al*., 2017), we examined how synchronized informative, uninformative, and random auditory stimuli paired with visual stimuli altered learning rate compared to a visual-stimuli only condition. In our last study, we contrasted rates of learning for two bimodal sequences where the auditory stimuli were either informative or uninformative.

Study 1 showed that sequence learning with bimodal stimuli shows a hallmark feature of traditional sequence learning tasks: sensitivity to the response-stimulus interval. In traditional unimodal sequence learning tasks, response-stimulus intervals of 500 ms improves sequence learning compared to no response-stimulus interval (Destrebecqz & Cleeremans, 2001), or in comparison to briefer response-stimulus intervals (Witt *et al*., 2023). Similarly, we also observe this pattern of results with bimodal stimuli, reflecting the impact of the task’s temporal dynamics on learning rates. Specifically, we found an initial advantage in speeding up reaction times for correct keypresses when the time between stimuli was longer (500 ms vs, 200 ms in Block 2, see Figure 2A). This initial advantage was concentrated at the early stages of learning, it was no longer prominent by the fourth block of trials, such that there was no difference between conditions in the final block of trials. The analysis of error rates revealed that participants made more errors as the block progressed, but they occurred more frequently when the response-stimulus interval was shorter across all blocks of trials. Thus, the interval immediately after a motor response is important not only to boost the rate of learning but also to reduce the number of errors in these types of tasks. These findings are consistent with the idea that after a motor response is made, it is beneficial to allow an assessment of the response to detect error (Huang & Shadmehr, 2007), and this error processing appears to involve the primary motor cortex Hadipour-Niktarash *et al*. (2007). One possibility is that by increasing the response-stimulus interval, we increased the opportunity for sensory predictions to take place and guide the advance preparation of motor actions, resulting in fewer mistakes overall.

In our second study, we showed that informative (congruent) audio-visual stimuli may confer learning benefits in comparison to non-informative (fixed or random) audio-visual stimuli, although we note that the difference was not statistically significant after corrections for multiple comparisons under the frequentist approach. In contrast to Silva *et al*. (2017), who found random auditory stimuli to be beneficial compared to visual only and equivalent to informative auditory stimuli during learning, we found that random auditory stimuli reached the final block of trials with an equivalent performance to the visual only group, but worse performance than the group who received informative auditory stimuli: Bayesian analysis supported this contrast, with a 95% credible interval of [−0.0545, −0.0233] and a posterior probability of direction (PD) = 0.0000. Therefore, while informative auditory stimuli did not consistently improve learning across all blocks, our findings suggest that their benefits may emerge in later blocks. These findings are consistent with those reported by Han *et al*. (2023) who found beneficial effects of synchronized informative auditory stimuli using a SISL task.

The results of Study 2 indicate that informative auditory stimuli may facilitate the rate of learning, particularly in later blocks, in comparison to random and no auditory stimuli. While differences between the informative and non-informative (single-tone) groups were not significant in the frequentist analysis, Bayesian analysis provided some support for this contrast in later blocks. This pattern suggests that the benefits of informative stimuli may emerge gradually rather than producing an immediate advantage. Given the potential influence of between-group variability, in our final study, we required participants to learn two sequences of actions: one paired with informative auditory stimuli and the other with uninformative (single-tone) stimuli. This within-subjects design allowed for a more controlled comparison, revealing that informative auditory stimuli resulted in greater training-related reductions in reaction times, starting from the second block of training trials.

One potential explanation for the difference between the informative and non-informative (single tone) groups in Study 2 and 3 is through reduced arousal from boredom associated with the single-tone auditory stimuli. However, this proposal is not wholly supported by previous evidence. For example, in a sequence learning task where tones are used as auditory feedback, similar to our findings, reaction times are faster with informative tone feedback compared to with non-informative single tone feedback, and compared to uninformative, random tone feedback (Keller & Koch, 2008). A boredom effect seems unlikely to wholly explain the faster reaction times in this previous work, because reaction times with non-informative single-tone feedback was similar to random uninformative tone feedback (Keller & Koch, 2008), which should, if anything, reduce boredom due to the lack of predictability of tones. Thus, based on this work and our current findings, we suggest that the benefit from auditory cues here result from the predictive content of the auditory cues, improving predictions about the motor sequence. Nonetheless, future experiments which explicitly quantify state arousal (e.g., via pupillometry) might help fully investigate the possibility that boredom underpins the slower reaction times shown with uninformative, single-tone auditory cues.

It is noteworthy that not all types of multisensory stimuli are effective in enhancing sequence learning. For example, pairing tactile stimuli with visual stimuli failed to alter sequence learning (Abrahamse *et al*., 2009). It is unclear if this beneficial effect of bimodal sequence learning is specific to the auditory domain. One possibility is that the close relationships between the auditory and motor areas facilitates learning the relationships between auditory stimuli and motor actions. For example, a large body of work (Chen *et al*., 2006; Zatorre *et al*., 2007; Chen *et al*., 2008; 2009; Grahn & Rowe, 2009; Lega *et al*., 2016) demonstrates the critical interplay between auditory and motor areas for musical performance. These associations can be learned quickly and are active not only when actions are required, but also when people simply listen to the sequence they were trained on (Lahav *et al*., 2007), suggesting an automatic engagement of auditory and motor areas in a similar context to ours.

An alternative, not mutually exclusive explanation is that informative auditory tonal stimuli used here might have altered intrinsic motivational and attentional factors known to affect motor learning (Wulf & Lewthwaite, 2016). Participants may have preferred the informative sequence over the uninformative, devoting more attention to it and, consequently, enhanced performance in that condition. Therefore, while auditory-motor networks might be automatically engaged when audio-visual stimuli are presented (Lahav *et al*., 2007), higher level cognitive processes might interact with these networks, prioritizing certain sensory inputs, such as auditory, when they are of intrinsic interest to the individual. Future experiments that explicitly manipulate such motivational and attentional factors (for example, by using informative and uninformative auditory stimuli that sensitively modulate intrinsic motivation/attention in participants) might help dissociate the contributions of motivational/attentional factors to bimodal sequence learning. Additionally, future studies could measure physiological markers of engagement such as pupil dilation, which Bianco *et al*. (2019) found was enhanced for both predictable and liked stimuli. This might help quantify the relative contributions of predictability and motivational factors in driving learning outcomes.

### 4.1. Limitations and future directions

In the work presented here, we focus our analyses on the learning phase of skill acquisition, seeking to determine whether accessory auditory stimuli could increase the rate of learning during practice. We compared our results against equivalent experimental phases in Silva *et al*. (2017) and Han *et al*. (2023) work. However, we did not test learning effects after a consolidation period nor without the presence of the auditory stimuli, except for the visual only group in study 2. Therefore, it remains to be tested if the beneficial effects for informative auditory stimuli on sequence learning remain after a period of consolidation (Walker *et al*., 2003) and when the auditory stimulus is removed during this later recall. It seems to us that without explicit knowledge, there is little evidence for a benefit of bimodal practice (Silva *et al*., 2017). This brings us to another important difference between our work and previous research in the literature, we clearly informed our participants they would be practicing sequences of actions and that knowing the sequence would improve performance. It is well established that explicit knowledge enhances performance in sequence learning tasks (Wong *et al*., 2015), so our results may only generalize to conditions where explicit learning is occurring. Future research should explore whether the factors we found to be important for learning in our task —time between stimuli and information contained in the auditory stimuli— impact implicit learning.

Another promising avenue for future research is to examine whether different forms of informative auditory stimuli improve sequence learning. For example, previous work suggests that pitch height has an inherent spatial mapping, where high pitches are associated with “up” and “right” responses while low pitches are associated with “down” and “left” responses (the SMARC effect) (Rusconi *et al*., 2006), and motor planning and motor performance is often improved by congruent pitch-spatial mappings. For example, in a sequence execution task, reaction times are faster when the spatial location of visual stimuli is congruent with the pitch of auditory feedback (Keller & Koch, 2008). Experience enhances effects of this pitch-spatial congruence, such that experienced musicians are less flexible than non-musicians when faced with inverted pitch-spatial mappings (Pfordresher & Chow, 2019). It is noteworthy however that the majority of previous work have examined effects of pitch-spatial congruence between auditory *feedback* and visual cues on performance of explicitly learnt sequences. To the best of our knowledge, no previous studies have examined the question of how pitch-spatial congruence between auditory and visual *cues* might alter sequence learning, as quantified in the conventional SRTT task used here. In our study, it is possible that pitch-spatial congruence in the informative condition might have driven performance benefits compared to the uninformative condition. To test this hypothesis, future studies can use a control condition that differentiates the distinct auditory cues via other stimuli properties such as timbre. Alternatively, auditory cues might be distinguished via loudness, which seem to have a spatial mapping that is independent from pitch (Koch *et al*., 2024).

### 4.2. Conclusion

In conclusion, our results demonstrate that the interval between each response and the subsequent stimuli in a bimodal SRT task can impact learning. They also indicate that informative auditory stimuli improve performance during learning. We speculate that there is a complex interplay between the temporal dynamics of the task, sensorimotor cortical networks, and attentional mechanisms mediating these effects. Although more research is needed to fully understand how accessory stimuli might benefit motor learning, our findings point to potential applications in applied contexts such as education and rehabilitation, where tailored multisensory interventions could enhance learning and skill acquisition.

## Acknowledgments

The authors would like to thank all study participants.

## Data availability statement

Data will be made available upon reasonable request.

## Author contributions

Li-Ann Leow: writing—review and editing, methodology. An Nguyen: Conceptualization, methodology, supervision of data collection. Emily Corti: methodology, supervision of data collection, writing—review. Welber Marinovic: Resources; conceptualization; methodology; investigation; formal analysis; writing—original draft.

## References

Abrahamse, E.L., van der Lubbe, R.H. & Verwey, W.B. (2009) Sensory information in perceptual-motor sequence learning: visual and/or tactile stimuli. Experimental brain research, 197, 175–183.

Baker, J.M. & Jordan, K.E. (2015) Chapter 11 - The Influence of Multisensory Cues on Representation of Quantity in Children. In Geary, D.C., Berch, D.B., Koepke, K.M. (eds) Mathematical Cognition and Learning. Elsevier, pp. 277–301.

Barr, D.J., Levy, R., Scheepers, C. & Tily, H.J. (2013) Random effects structure for confirmatory hypothesis testing: Keep it maximal. Journal of memory and language, 68, 255–278.

Berry, D.A. & Hochberg, Y. (1999) Bayesian perspectives on multiple comparisons. Journal of Statistical Planning and Inference, 82, 215–227.

Bianco, R., Gold, B., Johnson, A. & Penhune, V. (2019) Music predictability and liking enhance pupil dilation and promote motor learning in non-musicians. Scientific reports, 9, 17060.

Chen, J.L., Penhune, V.B. & Zatorre, R.J. (2008) Listening to musical rhythms recruits motor regions of the brain. Cerebral Cortex, 18, 2844–2854.

Chen, J.L., Penhune, V.B. & Zatorre, R.J. (2009) The role of auditory and premotor cortex in sensorimotor transformations. Annals of the New York Academy of Sciences, 1169, 15–34.

Chen, J.L., Zatorre, R.J. & Penhune, V.B. (2006) Interactions between auditory and dorsal premotor cortex during synchronization to musical rhythms. Neuroimage, 32, 1771–1781.

Cohen, A., Ivry, R.I. & Keele, S.W. (1990) Attention and structure in sequence learning. *Journal of Experimental Psychology: Learning*, Memory, and Cognition, 16, 17.

Curran, T. & Keele, S.W. (1993) Attentional and Nonattentional Forms of Sequence Learning. *Journal of Experimental Psychology: Learning*, Memory, and Cognition, 19, 189–202.

Destrebecqz, A. & Cleeremans, A. (2001) Can sequence learning be implicit? New evidence with the process dissociation procedure. Psychonomic bulletin & review, 8, 343–350.

Flores, R.D., Sanders, C.A., Duan, S.X., Bishop-Chrzanowski, B.M., Oyler, D.L., Shim, H., Clocksin, H.E., Miller, A.P. & Merkle, E.C. (2022) Before/after Bayes: A comparison of frequentist and Bayesian mixed-effects models in applied psychological research. Br J Psychol, 113, 1164–1194.

Grahn, J.A. & Rowe, J.B. (2009) Feeling the beat: premotor and striatal interactions in musicians and nonmusicians during beat perception. J Neurosci, 29, 7540–7548.

Hadipour-Niktarash, A., Lee, C.K., Desmond, J.E. & Shadmehr, R. (2007) Impairment of retention but not acquisition of a visuomotor skill through time-dependent disruption of primary motor cortex. The Journal of neuroscience : the official journal of the Society for Neuroscience, 27, 13413–13419.

Han, Z., Sanchez, D., Levitan, C.A. & Sherman, A. (2023) Stimulus-locked auditory information facilitates real-time visuo-motor sequence learning. Psychonomic Bulletin & Review.

Hedeker, D. & Gibbons, R.D. (2006) Longitudinal data analysis. Wiley-Interscience.

Hoffmann, J. & Koch, I. (1997) Stimulus-response compatibility and sequential learning in the serial reaction time task. Psychological Research, 60, 87–97.

Huang, V.S. & Shadmehr, R. (2007) Evolution of motor memory during the seconds after observation of motor error. Journal of neurophysiology, 97, 3976–3985.

Keller, P.E. & Koch, I. (2008) Action planning in sequential skills: Relations to music performance. Quarterly Journal of Experimental Psychology, 61, 275–291.

Kim, R.S., Seitz, A.R. & Shams, L. (2008) Benefits of stimulus congruency for multisensory facilitation of visual learning. PLoS One, 3, e1532.

Koch, S., Schubert, T. & Blankenberger, S. (2024) Simultaneous but independent spatial associations for pitch and loudness. Psychological Research, 1–14.

Kuznetsova, A., Brockhoff, P.B. & Christensen, R.H.B. (2015) Package ‘lmertest’. R package version, 2, 734.

Lahav, A., Saltzman, E. & Schlaug, G. (2007) Action representation of sound: audiomotor recognition network while listening to newly acquired actions. J Neurosci, 27, 308–314.

Lega, C., Stephan, M.A., Zatorre, R.J. & Penhune, V. (2016) Testing the Role of Dorsal Premotor Cortex in Auditory-Motor Association Learning Using Transcranical Magnetic Stimulation (TMS). PLoS One, 11, e0163380.

Luan, M., Maurer, H., Mirifar, A., Beckmann, J. & Ehrlenspiel, F. (2021) Multisensory action effects facilitate the performance of motor sequences. Atten Percept Psychophys, 83, 475–483.

Makowski, D., Ben-Shachar, M.S. & Lüdecke, D. (2019) bayestestR: Describing effects and their uncertainty, existence and significance within the Bayesian framework. Journal of Open Source Software, 4, 1541.

Morey, R.D. (2008) Confidence intervals from normalized data: A correction to Cousineau (2005). Tutorials in Quantitative Methods for Psychology, 4, 61–64.

Morey, R.D., Rouder, J.N., Jamil, T. & Morey, M.R.D. (2015) Package ‘bayesfactor’. URLh http://cran/r-projectorg/web/packages/BayesFactor/BayesFactor *pdf i (accessed 1006 15)*.

Nissen, M.J. & Bullemer, P. (1987) Attentional requirements of learning: Evidence from performance measures. Cognitive Psychology, 19, 1–32.

Perez, M.A., Wise, S.P., Willingham, D.T. & Cohen, L.G. (2007) Neurophysiological mechanisms involved in transfer of procedural knowledge. Journal of Neuroscience, 27, 1045–1053.

Pfordresher, P.Q. & Chow, K. (2019) A cost of musical training? Sensorimotor flexibility in musical sequence learning. Psychonomic Bulletin & Review, 26, 967–973.

Robinson, C.W. & Parker, J.L. (2021) Tones slow down visuomotor responses in a visual-spatial task. Acta psychologica, 218, 103336.

Rusconi, E., Kwan, B., Giordano, B.L., Umilta, C. & Butterworth, B. (2006) Spatial representation of pitch height: the SMARC effect. Cognition, 99, 113–129.

Seitz, A.R., Kim, R. & Shams, L. (2006a) Sound facilitates visual learning. Current Biology, 16, 1422–1427.

Seitz, A.R., Kim, R. & Shams, L. (2006b) Sound facilitates visual learning. Curr Biol, 16, 1422–1427.

Shams, L. & Seitz, A.R. (2008) Benefits of multisensory learning. Trends Cogn Sci, 12, 411–417.

Silva, A.E., Barakat, B.K., Jimenez, L.O. & Shams, L. (2017) Multisensory Congruency Enhances Explicit Awareness in a Sequence Learning Task. Multisensory Research, 30, 681–689.

Stark-Inbar, A., Raza, M., Taylor, J.A. & Ivry, R.B. (2017) Individual differences in implicit motor learning: task specificity in sensorimotor adaptation and sequence learning. Journal of neurophysiology, 117, 412–428.

Stocker, C. & Hoffmann, J. (2004) The ideomotor principle and motor sequence acquisition: tone effects facilitate movement chunking. Psychol Res, 68, 126–137.

Stocker, C., Sebald, A. & Hoffmann, J. (2003) The influence of response--effect compatibility in a serial reaction time task. Q J Exp Psychol A, 56, 685–703.

van Vugt, F.T. & Tillmann, B. (2015) Auditory feedback in error-based learning of motor regularity. Brain research, 1606, 54–67.

Vigliocco, G., Perniss, P. & Vinson, D. (2014) Language as a multimodal phenomenon: implications for language learning, processing and evolution. Philos Trans R Soc Lond B Biol Sci, 369, 20130292.

Walker, M.P., Brakefield, T., Allan Hobson, J. & Stickgold, R. (2003) Dissociable stages of human memory consolidation and reconsolidation. Nature, 425, 616–620.

Willingham, D.B., Greenberg, A.R. & Thomas, R.C. (1997) Response-to-stimulus interval does not affect implicit motor sequence learning, but does affect performance. Memory & Cognition, 25, 534–542.

Witt, A., Poulin-Charronnat, B., Bard, P. & Vinter, A. (2023) The effect of response-to-stimulus interval on children’s implicit sequence learning. Journal of Experimental Child Psychology, 232, 105668.

Wong, A.L., Lindquist, M.A., Haith, A.M. & Krakauer, J.W. (2015) Explicit knowledge enhances motor vigor and performance: motivation versus practice in sequence tasks. J Neurophysiol, 114, 219–232.

Wulf, G. & Lewthwaite, R. (2016) Optimizing performance through intrinsic motivation and attention for learning: The OPTIMAL theory of motor learning. Psychonomic Bulletin & Review, 23, 1382–1414.

Yu, Z., Guindani, M., Grieco, S.F., Chen, L., Holmes, T.C. & Xu, X. (2022) Beyond t test and ANOVA: applications of mixed-effects models for more rigorous statistical analysis in neuroscience research. Neuron, 110, 21–35.

Zatorre, R.J., Chen, J.L. & Penhune, V.B. (2007) When the brain plays music: auditory-motor interactions in music perception and production. Nature Reviews. Neuroscience, 8, 547–558.

